# Vulnerable caribou and moose populations display contrasting responses to mountain pine beetle outbreaks and management

**DOI:** 10.1101/2024.09.23.614365

**Authors:** Laura L. Griffin, Laura Finnegan, Julie Duval, Simone Ciuti, Virginia Morera-Pujol, Haitao Li, A. Cole Burton

## Abstract

1. Rising global temperatures and changing landscape conditions have led to widespread mountain pine beetle (MPB) outbreaks in western North America. Extensive management has been implemented in response, via forest harvesting and prescribed burning. However, the impacts of MPB and MPB-management on ungulate populations, particularly caribou and moose, remain poorly studied. Given the differing specialisations of these species (caribou are habitat specialists, moose are generalists), their responses to these disturbances may vary. This could, in turn, lead to unintended, indirect effects (e.g., disturbance-mediated apparent competition, whereby increased moose presence results in increased predator presence, with resultant caribou declines). Unravelling these responses is crucial for informing MPB management and ensuring that applied actions do not exacerbate adverse effects.
2. We assessed the effects of early-stage MPB-infestation, harvest, and fire, on habitat selection by caribou (boreal and central mountain designatable units) and moose in west-central Alberta. We built resource selection functions and functional response models using GPS collar data collected 3-5 years after MPB infestation and found that responses varied between species.
3. Caribou had complex seasonal responses to MPB, generally avoiding areas with more MPB disturbance in winter but using them in summer. Caribou typically avoided harvested and burned areas, though this was dependent on the overall degree of disturbance within their ranges. In contrast, moose had positive responses to both MPB and burned areas year-round.
4. These findings suggest that MPB may negatively impact long-term winter habitat availability for threatened caribou populations, but may have positive impacts on moose habitat. However, moose use of MPB-impacted areas could further affect caribou by contributing to disturbance-mediated apparent competition.
5. *Synthesis and applications:* The negative impacts of forest harvest and burning on caribou suggests that less intensive actions are required for MPB management within caribou ranges.. While moose selectively used harvested and burned areas, it is possible that they may have adverse responses to cumulative disturbance over time. As caribou and moose responded to MPB soon after infestation, ongoing monitoring is required to detect MPB early and facilitate proactive management, though further study is needed to determine the most effective actions.

## 1. Introduction

Global biodiversity loss and wildlife population declines have prompted research into how different disturbance types affect natural systems (Brodie, Williams and Garner, 2021; Pirotta *et al*., 2018). Understanding how emerging disturbances impact newly exposed wildlife populations enables landscape managers to reduce and mitigate the negative effects of disturbances on vulnerable species (Northrup and Wittemyer, 2013). One emerging disturbance gaining significant attention is insect outbreaks, which have become progressively more common with climate change and forest homogenization (Ayres and Lombardero, 2018; Ono, 2004).

Mountain pine beetle (MPB; *Dendroctonus ponderosae*) is a bark beetle native to the pine forests of western North America. It periodically erupts into large-scale outbreaks, drastically altering forest ecosystems (Mitton and Ferrenberg, 2012; Sambaraju and Goodsman, 2021). MPB typically targets large, mature pines, reducing canopy cover and decreasing shade-tolerant understory species while shade-intolerant vascular plant species increase (Pec *et al*., 2015; Schoennagel *et al*., 2012; Steinke *et al*., 2020). In its most recent outbreak, MPB has spread to higher latitudes and elevations than previously documented and beyond its historic range, including into western Alberta, Canada (Howe *et al*., 2021; Sambaraju and Goodsman, 2021). There is significant concern about the impacts of MPB within these novel areas, including how local wildlife species are responding to this new disturbance (Ono, 2004; Saab *et al*., 2014).

In general, the impacts of MPB on wildlife are understudied, with predictions often extrapolated from studies assessing vegetation changes (Chan-McLeod, 2006). The few existing assessments of wildlife responses to MPB have primarily focused on avian species, and there is a scarcity of literature on the responses of large mammals, despite their cultural, ecological, and economic importance (Festa-Bianchet *et al*., 2011; Popp *et al*., 2020; Saab *et al*., 2014). This limits the application of informed MPB management policies aiming to reduce negative effects on large mammalian species, particularly in newly impacted areas (Dhar, Parrott and Heckbert, 2016). Instead, blanket management practices, including accelerated pine harvest, salvage logging, and prescribed burns, have been applied (ASRD, 2007; McClelland *et al*., 2023). However, these actions could cause more adverse responses in vulnerable mammal populations than MPB itself, as they change understory vegetation structure and composition (Steinke *et al*., 2020). To inform effective landscape management, we must, therefore, unravel how mammals respond to both MPB and MPB management.

Large ungulates play a key role in forest ecosystems and may be directly impacted by changes to understory forage availability (Boan, McLaren and Malcolm, 2011). In western Canada, how forest disturbance impacts woodland caribou (*Rangifer tarandus caribou*) is of particular interest as caribou are federally and provincially listed as Threatened (Government of Alberta, 2022; Government of Canada, 2018). Moose (*Alces alces*) are also of interest as populations are in general decline across many parts of Canada, but increasing in select areas within western Canada (Government of Alberta, 2012-2024.; Kuzyk *et al*., 2018). Moose are integral to Indigenous food security, resident hunting opportunities (Priadka *et al*., 2022), and are an apparent competitor of caribou (Neufeld *et al*., 2021). It has been theorised that moose population fluctuations may be linked to changing environmental factors and landscapes (Anderson, McLellan and Serrouya, 2018; Neufeld *et al*., 2021), highlighting the need for research to determine how moose populations are affected by forest changes and management practices.

Caribou and moose have different habitat requirements, so they may respond differently to MPB and MPB management (Belovsky, 1981; Webber *et al*., 2022). Caribou are habitat specialists; they are dietarily reliant on lichens in mature forests during winter, though they also eat vascular plants during summer (Nobert *et al*., 2020; Webber *et al*., 2022). MPB thins canopy cover, causing declines in lichen abundance as early as 3-5 years post-infestation, though declines may not occur for 10-15 years depending on local environmental factors (Cichowski *et al*., 2022; Lewis and Hartley, 2006b; Nobert *et al*., 2020). MPB-killed trees may also begin to fall 3-5 years post-infestation, potentially impeding ungulate movement (van Ginkel *et al*., 2021). It is, therefore, theorised that caribou may be negatively impacted by MPB infestations, but this has not been assessed (Cichowski *et al*., 2022). Conversely, while lichen cover may decrease soon after MPB infestation, the abundance and diversity of other understory vegetation generally increases (Cichowski *et al*., 2022; Seip and Jones, 2008; Stone and Wolfe, 1996). Moose are generalist herbivores, feeding on a wider range of plant species year-round than caribou (Belovsky, 1981). As MPB increases understory plant growth, MPB may have positive impacts on moose, but this also remains untested (Pappas, Tinker and Rocca, 2020; Steinke *et al*., 2020).

In terms of MPB management, caribou are sensitive to habitat disturbance, and avoid recently harvested, salvage logged, or burned areas (Finnegan *et al*., 2021; Seip and Jones, 2008). In contrast, moose typically select recently harvested or burned areas, once early seral vegetation is regrowing (Johnson and Rea, 2023a; Wasser *et al*., 2011). In some cases, this has resulted in disturbance-mediated apparent competition (DMAC), where disturbance-caused habitat changes result in increased abundances of moose and/or other generalist ungulates (e.g., deer) and, subsequently, increased abundances of shared predators. This, therefore, increases predation risk for caribou and drives them out of these areas (Environment and Climate Change Canada, 2018; Neufeld *et al*., 2021). However, this may be context dependent (DeMars *et al*., 2019; Mumma *et al*., 2018) and moose declines have also been documented in areas with increased habitat disturbance (Koetke *et al*., 2023). Determining how caribou and moose respond to MPB and MPB management can, therefore, inform actions to mitigate the impacts of landscape management on both species, while proactively guiding effective stewardship of forest ecosystems.

In this study, we used GPS collar data from caribou and moose to assess responses to MPB and MPB management. For MPB, we focused on the early period of MPB infestation (3–5 years old), when changes to forest and vegetation structure first occur, and when initial management responses may be implemented. As a proxy for MPB management, we focused on harvest blocks and wildfires < 20 years old, when disturbed patches are dominated by vascular plants (Finnegan *et al*., 2021; Lacerte, Leblond and St Laurent, 2021; Silva, 2020). We predicted that moose would use habitats disturbed by MPB and associated management practices, whereas caribou would avoid them.

Specifically, we predicted that:

1. As mature forest specialists, caribou would avoid areas with:

a. MPB (3–5 years old) due to associated impacts on understory vegetation (i.e., loss of lichens, increase of other vegetation species) and because caribou generally avoid disturbed forest.
b. Recent harvest and fires (≤20 years old) because caribou generally avoid disturbance (i.e., due to loss of forage and increased risk of predation in these areas).
c. We also predicted that caribou would have a functional response to disturbance, meaning that the strength of avoidance of MPB, harvest, and fire increases as that disturbance becomes more prevalent within an individual’s home range.
2. Moose, as generalists:

a. Would elect areas with more disturbance (MPB and MPB management).
b. Would not have a functional response to disturbance, instead selecting disturbed habitat relative to availability within their home ranges.

## 2. Methods

### 2.1 Study area

The study area was in west-central Alberta, Canada (Fig 1), one of the initial entry points of MPB into the area east of the Rocky Mountains, with MPB identified via government helicopter surveys in 2005 (Government of Alberta and Tyssen, 2009). We focused on the area encompassing the range of two populations of woodland caribou; one boreal (Little Smoky-LS) and one central mountain (Redrock-Prairie Creek-RPC) (Fig. 1) (Hervieux *et al*., 2013). The focal moose population overlaps with these caribou ranges (Fig. 1). This area was within the traditional territories, meeting grounds, and homes of many Indigenous Peoples, including the the Aseniwuche Winewak, Danezaa, Métis, Nehiyawak, Simpcw, Stoney and Tsuut’ina (best available information from Native-land.ca).

**Fig. 1.**
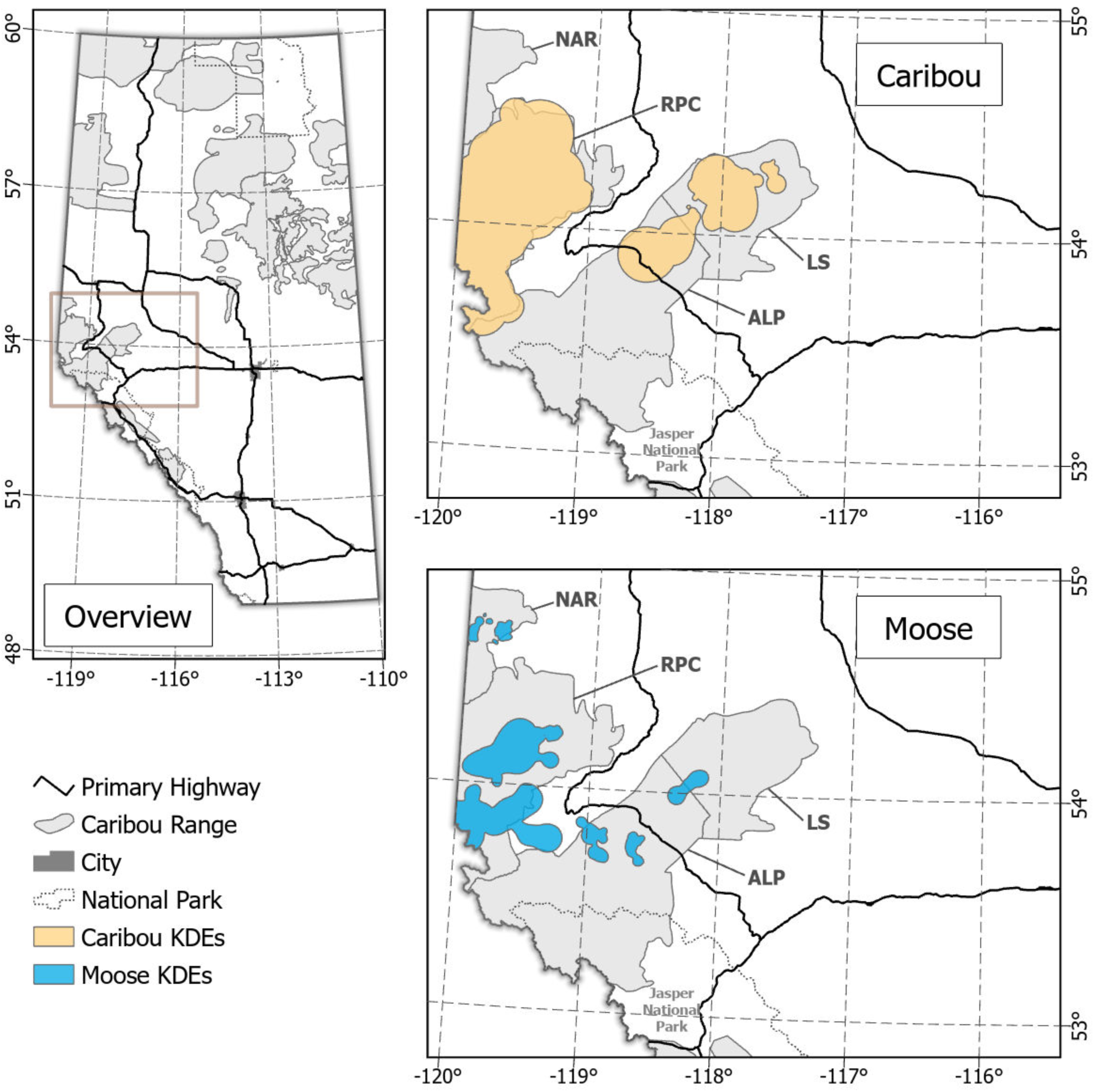
Caribou (orange) and moose (blue) home ranges in west-central Alberta, Canada, used to assess responses to mountain pine beetle between 2008 and 2010. Home ranges are shown on a map of the overall province of Alberta (left), with the zoomed-in inlay (left) represented by a grey square. Population ranges for the A La Peche (ALP), Little Smoky (LS), Narraway (NAR), and Redrock-Prairie Creek (RPC) caribou are outlined and labelled.

The study area includes alpine, subalpine, upper foothills, and lower foothills natural subregions. It is characterised by coniferous forests with lodgepole pine (*Pinus contorta*), being the primary host for MPB in these habitats (Dempster and Meredith, 2021; Natural Regions Committee, 2006). Treed muskegs are also present, primarily in the Little Smoky range. As well as disturbance from MPB, there is anthropogenic disturbance, including forest harvesting and oil and gas extraction, and natural disturbance, including wildfire and windthrow. Further details on the area is available in McClelland et al. (2023).

### 2.2 Animal location data

We obtained GPS location data from female caribou collared by the Government of Alberta as part of long-term provincial monitoring programs. Capture and collaring is described in more detail in Hervieux et al. (2013), and collars collected location information with a 1.5-4h fix rate (Lotek GPS1000, 2000, 2200, 3300, 4400 models; Lotek Engineering, Newmarket, Ontario, Canada). We kept boreal and central mountain caribou designatable units separate for analyses (Weckworth *et al*., 2018). Hereafter, whenever we refer to ‘boreal caribou’ we are specifically referring to the Little Smoky population and whenever we refer to ‘central mountain caribou’ we are referring to the Redrock-Prairie Creek population. We acknowledge that the results extracted from these populations may not be transferable to other populations of the same designatable unit, and do not wish to overextend interpretations, but have abbreviated to the designatable unit in places (e.g., figures and tables) to improve readability.

The focal moose population overlaps with these caribou ranges and extends into the Narraway and A La Peche central mountain caribou ranges (Fig. 1). Male and female moose GPS collar data were originally collected as part of a study on moose habitat in west-central Alberta, and were obtained from Movebank for this study (2-4h fixes; ATS G2000 GPS collars, Advanced Telemetry Systems, Isanti, MN, USA) (www.movebank.org, Peters *et al*., 2013).

Data handling and analyses were performed in R version 4.3.2 (R Core Team, 2023). Location data for both species were collected from 2008-2010, 3–5 years after MPB infestations first occurred. We removed locations with high dilution of precision values (DOPs: >12 for caribou and >5 for moose) or that occurred outside of a 2-4 hour fix interval. Caribou data were pre-cleaned to a DOP >12, and further information on DOP and number of satellites were not provided. For moose data, we removed 2D points and points with a DOP >5 (sensu Lewis *et al*., 2007). We also removed individual caribou and moose with a fix-rate success of <90% (sensu Frair *et al*., 2010; Hebblewhite, Percy and Merrill, 2007), and removed locations with an incoming and outgoing speed above the 99^th^ percentile (Gupte *et al*., 2022). Further details on data cleaning are in S1.

For each dataset, we then partitioned locations by individual, year, and season: winter (October 16^th^ to May 15^th^) and summer (May 16^th^ to October 15^th^) (Peters *et al*., 2013). Hereafter, individual-year refers to individual-season-year combinations. We generated annual seasonal home ranges for each individual-year using kernel density estimation (95% adaptive kernel, default bandwidth) in *amt* (Signer, Fieberg and Avgar, 2019). We clipped home ranges and location data to exclude British Columbia and Jasper National Park, where beetle data were not available. We removed individual-year combinations without MPB in their home range and removed individual-year combinations with <200 locations (S1&2). We generated 10 available locations for each used location within seasonal individual-year home ranges via *amt* (Signer, Fieberg and Avgar, 2019). The final boreal caribou dataset included locations from 16 individual-years (n=9 winter, n=7 summer, 5 collared animals) (Fig. 1&S2). The final central mountain caribou dataset included locations from 45 individual-years (n=32 winter, n=13 summer, 20 animals) (Fig. 1&S2). The final moose dataset included locations from 24 individual-years (n=16 winter, n=8 summer, 9 animals) (Fig. 1&S2).

### 2.3 Predictor variables

We performed geoprocessing and extraction of predictor variables in ArcGIS Pro 3.1.2 (ESRI, 2023), and QGIS 3.32.1 (QGIS Development Team, 2023). For MPB, we extracted annual MPB survey data available from the Government of Alberta (Government of Alberta, 2023). We combined these data to create annual 30×30m rasters, with each cell depicting whether that area had been disturbed by MPB or not (Fig. 2a, S3).

**Fig. 2.**
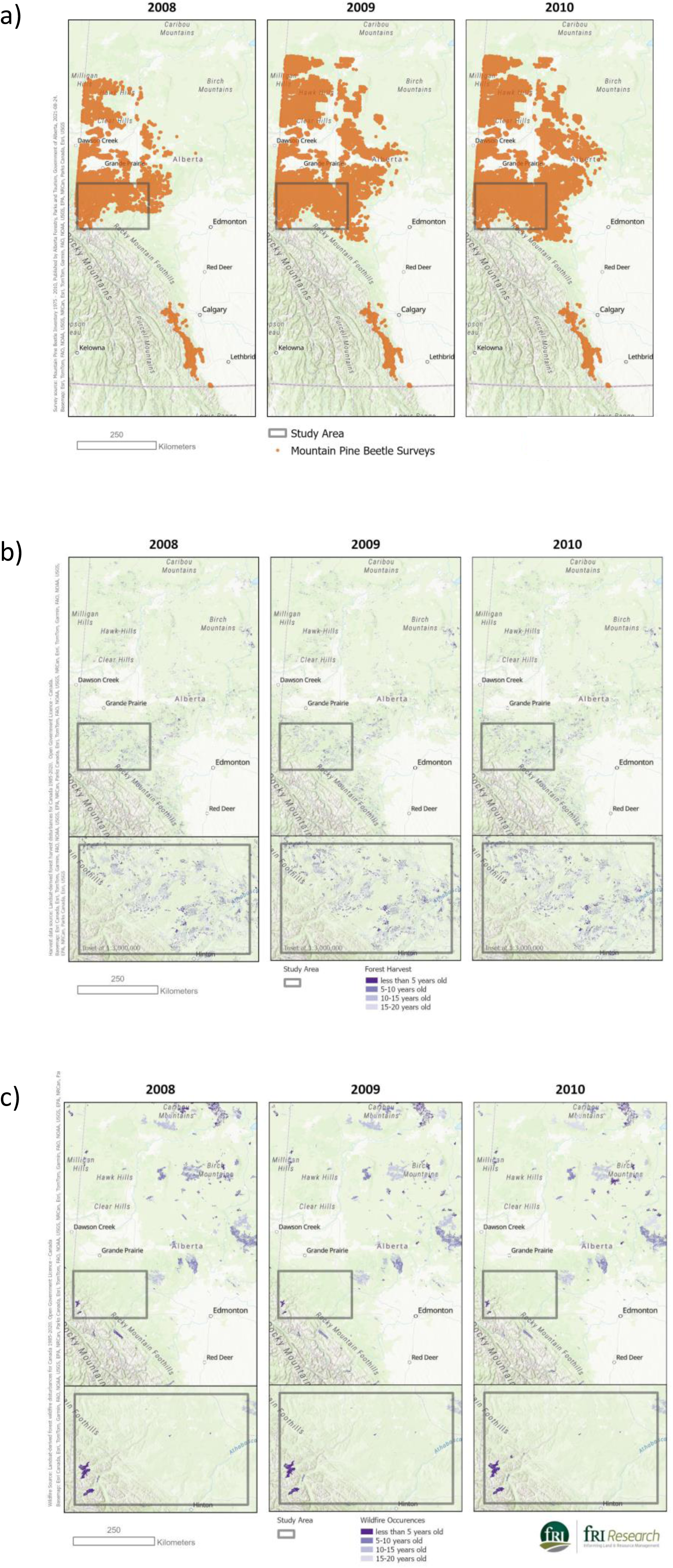
Area impacted by mountain pine beetle (MPB) (a), ≤20 year old harvest (b), and ≤20 year old fire (c) in west-central Alberta, Canada, over the years of our study (2008-2010). MPB point locations represent 30×30m areas where MPB was detected by annual aerial surveys, including locations from both that year and previous years, that had not been removed by fires or harvest. Disturbance polygon colour vary with time since disturbance. The grey square highlights our study area, one of the initial entry points of MPB into Alberta.

For harvest and fire, we used Landsat-derived 30×30m rasters available from the National Terrestrial Ecosystem Monitoring system (NTEMS) (Hermosilla *et al*., 2018; Hermosilla *et al*., 2016) (Fig. 2b and 2c). These were used as proxies for recent MPB management via harvesting or prescribed burning. We recognise that accelerated pine harvesting and salvage logging in response to MPB may be even more intensive than regular harvesting activities (e.g., percentage removal, size of harvest areas), but those data were not available across our study area. Previous studies have used clearcuts as proxies for salvage logging and, given that some policies have included greater percentage tree retention in MPB management, we opted to use all harvest activity (Peter and Bogdanski, 2010; Steinke *et al*., 2020; Thorn *et al*., 2018). We subset harvest and fire rasters to only include recent (≤20-year-old) disturbances, as previously explained, and then generated densities at 1km and 5km radii around each animal location (Finnegan *et al*., 2021; Lacerte, Leblond and St Laurent, 2021; Silva, 2020).

To account for other landscape characteristics that impact habitat use, we also included land cover (NTEMS), elevation, and slope (Digital Elevation Model; Natural Resources Canada, 2011). We re-categorised landcover into forested and open locations (S3).

### 2.4 RSF Models

Before fitting models, we screened variables for collinearity (|r_p_| < 0.7) (Dormann *et al*., 2013); grouping 1km density variables together and 5km density variables together (i.e., 12 collinearity tests, S4). Harvest density at the 5km scale was collinear (r=-0.7) with elevation in moose data during summer (and borderline in winter, r=-0.6). Therefore, we dropped harvest from moose RSF models, but retained it for caribou models.

We fit each RSF as a binomial (used/available) generalised linear mixed effect model (GLMM) in *lme4* (Bates *et al*., 2015), including animal ID-season-year as a random effect, and using the “bobyqa” optimiser (Hedlin and Franke, 2017; Muthukrishnan, Hansel Welch and Larkin, 2018). We log-transformed and scaled numerical variables to improve model convergence and included them as single and quadratic terms, to allow for non-linear patterns. We applied the same model structure to each dataset (with the exception of harvest which was dropped from moose models), including sex as a categorical variable in moose models. We built models using disturbance densities at the 1km and 5km scales for each of the 6 datasets (one dataset for each population and season, i.e., 12 models (S5)). We used Akaike Information Criterion (AIC) to identify the most parsimonious scale for models, i.e., retaining the scale with the lowest AIC value (Sakamoto, Ishiguro and Kitagawa, 1986) (S5).

We assessed model fit using R2 for mixed models (Nakagawa and Schielzeth, 2013) in *MumIn* (Barton, 2019). We used variograms to check for spatial autocorrelation in model residuals, with none being flagged (Roberts *et al*., 2017) (S6). We assessed model prediction using five-fold cross validation; partitioning data into 80% training data and 20% testing data (Boyce *et al*., 2002). We computed Spearman rank correlations (r_s_) between frequencies of area-adjusted cross-validation locations, and then calculated a rank for each cross-validated model. Better predictive performance is indicated by a strong positive correlation value, demonstrating that model predictions match actual observations (Smith *et al*., 2022).

### 2.5 Functional Response (FR) Models

We generated functional response (FR) models on the multiplicative scale, as recommended by Holbrook et al. (2019). This allowed us to assess how habitat use changed relative to availability across home ranges (Mysterud and Ims, 1998). The null hypothesis posits that habitat use is a constant multiplicative function of habitat availability, meaning that the log_e_-transformed ratio of use:availability remains constant, with deviations indicating a functional response in habitat selection (Holbrook *et al*., 2019).

We used base R (R Core Team, 2023) to fit linear, quadratic, and polynomial models for each disturbance variable (i.e., MPB, harvest, and fire) to each of the six datasets (excluding harvest from moose models) at the 5km scale We used log_e_-transformed mean disturbance density for each individual-year’s used locations as the response variable and log_e_-transformed mean disturbance density for each individual-year’s available locations as a fixed effect. We determined if linear, quadratic, or polynomial models best fit the data using likelihood ratio tests via *lmtest* (Zeileis and Hothorn, 2002) (S7). We exponentiated the predictions for visualisation and interpretation, and visualised results in *ggplot2* (Wickham, 2016). We considered the overall trend in the relationship and the associated confidence intervals in assessments.

## 3. Results

### Little Smoky (Boreal caribou)

RSF models indicated that boreal caribou from the Little Smoky population avoided areas with higher densities of MPB during winter but selected them during summer (Table 1, Fig. 3a&3d). These caribou selected areas with higher densities of harvest during winter, and avoided harvest during summer (Table 1, Fig. 3b&3e). There was no response by these caribou to fire during winter, but they selected areas with higher densities of fire during summer (Table 1, Fig. 3c&3f), though notably fire was rare overall within home ranges in the Little Smoky population range (S3). Generally, these boreal caribou selected forested habitat during winter and open areas during summer. They selected lower elevations during winter, higher elevations during summer, and flatter areas (lower slopes) year-round (Table 1). The boreal caribou winter model had fair fit (23.53%) and the summer model had moderate fit (51.81%) (S8). K-fold cross-validation demonstrated that both models had a high predictive performance, with a mean Spearman correlation of r_s_=0.97 for winter, and r_s_=0.96 for summer (Mukaka, 2012) (S8-S9).

**Fig. 3.**
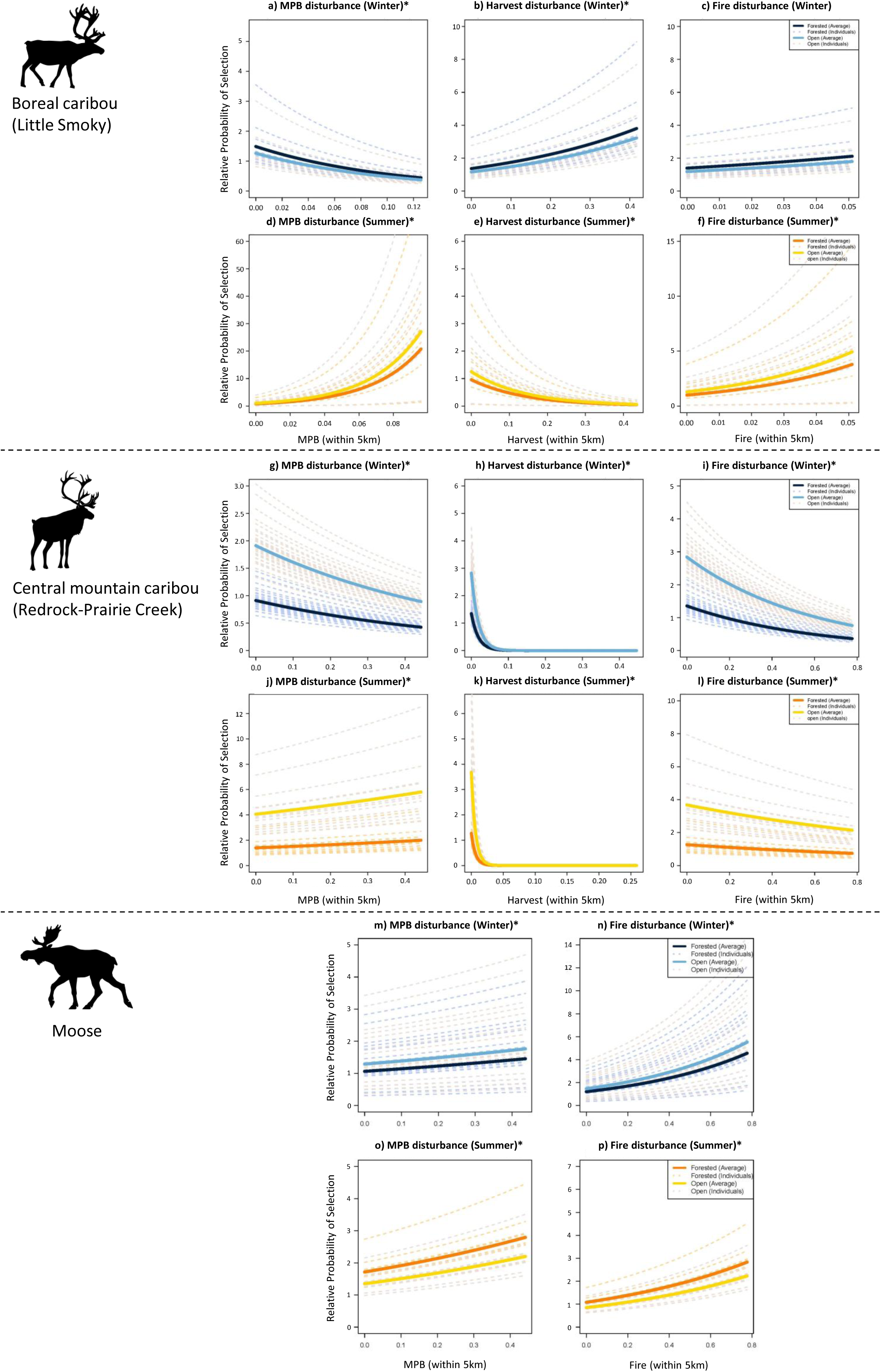
Relative probability of boreal caribou, central mountain caribou, and moose habitat selection in relation to MPB, harvest and fire during winter (blue) and summer (orange), in west-central Alberta, Canada, during 2008-2010. Harvest was excluded from moose models. The relative probability of selection is depicted when the individual is locally (30×30m) within forested or open habitat. All habitat variables represent % density at a 5km radius. Dashed lines are estimates for each individual caribou. Scales for relative probability of selection on the y-axis have been adjusted to improve legibility. Significant variables are marked with an asterisk.

**Table 1.** Estimated β coefficients, standard errors (SE), lower and upper 95% confidence intervals, Z-values, and p-values of boreal caribou habitat selection model during winter and summer in west-central Alberta, Canada, during 2008-2010. Reference category for habitat was ‘forested’.

| Fixed effects | Estimate | SE | Lower 95% CI | Upper 95% CI | Z-value | p-value |
| --- | --- | --- | --- | --- | --- | --- |
| **Boreal caribou: Winter** |  |  |  |  |  |  |
| Intercept | -2.347 | 0.128 | -2.598 | -2.096 | -18.326 | <0.001 |
| Habitat [Open] | -0.164 | 0.026 | -0.216 | -0.112 | -6.235 | <0.001 |
| Elevation (m) | 55.898 | 1.321 | 53.310 | 58.486 | 42.331 | <0.001 |
| Elevation<sup>2</sup> (m) | -56.592 | 1.327 | -59.194 | -53.991 | -42.638 | <0.001 |
| Slope | 0.105 | 0.0323 | 0.041 | 0.169 | 3.194 | 0.002 |
| Slope<sup>2</sup> | -0.434 | 0.036 | -0.505 | -0.363 | -12.013 | <0.001 |
| Harvest | 0.256 | 0.048 | 0.161 | 0.351 | 5.295 | <0.001 |
| Harvest<sup>2</sup> | 0.038 | 0.043 | -0.047 | 0.122 | 0.869 | 0.385 |
| Fire | 0.016 | 0.050 | -0.082 | 0.113 | 0.315 | 0.752 |
| Fire<sup>2</sup> | 0.059 | 0.0480 | -0.035 | 0.153 | 1.235 | 0.217 |
| MPB | -0.645 | 0.032 | -0.708 | -0.582 | -19.962 | <0.001 |
| MPB<sup>2</sup> | 0.286 | 0.030 | 0.227 | 0.344 | 9.605 | <0.001 |
| **Boreal caribou: Summer** |  |  |  |  |  |  |
| Intercept | -2.737 | 0.453 | -3.624 | -1.851 | -6.050 | <0.001 |
| Habitat [Open] | 0.265 | 0.032 | 0.202 | 0.328 | 8.233 | <0.001 |
| Elevation (m) | -18.360 | 2.523 | -23.306 | -13.414 | -7.276 | <0.001 |
| Elevation<sup>2</sup> (m) | 19.213 | 2.522 | 14.269 | 24.156 | 7.617 | <0.001 |
| Slope | -0.375 | 0.041 | -0.455 | -0.294 | -9.115 | <0.001 |
| Slope<sup>2</sup> | -0.087 | 0.049 | -0.183 | 0.010 | -1.763 | 0.078 |
| Harvest | 0.789 | 0.062 | 0.667 | 0.911 | 12.685 | <0.001 |
| Harvest<sup>2</sup> | -1.230 | 0.065 | -1.337 | -1.082 | -18.544 | <0.001 |
| Fire | -3.057 | 0.199 | -3.448 | -2.666 | -15.335 | **<0.001** |
| Fire<sup>2</sup> | 3.368 | 0.191 | 2.994 | 3.742 | 17.643 | **<0.001** |
| MPB | 0.717 | 0.090 | 0.541 | 0.894 | 7.983 | **<0.001** |
| MPB<sup>2</sup> | 0.067 | 0.076 | 0.083 | 0.217 | 0.879 | 0.380 |

FR models indicated that these boreal caribou did not exhibit a functional response to MPB during winter or summer, though there was a nonsignificant trend indicating avoidance of MPB in home ranges with higher MPB densities during winter (i.e., use less than proportional to availability) (Fig. 4a&4d). During winter, these caribou avoided harvest when in home ranges with lower harvest densities, but were more likely to use harvest in home ranges with higher harvest densities (Fig. 4b). During summer, their response to harvesting was proportional when in home ranges with lower densities of harvest, however, they were more likely to avoid it as densities increased across home ranges (Fig. 4e). There was no functional response to fire during winter, but during summer they were more likely to use fire as densities increased across home ranges (Fig. 4c&4f). While these significant non-linear effects indicate a functional response, the limited sample size resulted in large confidence intervals, limiting the strength of inferences.

**Fig. 4.**
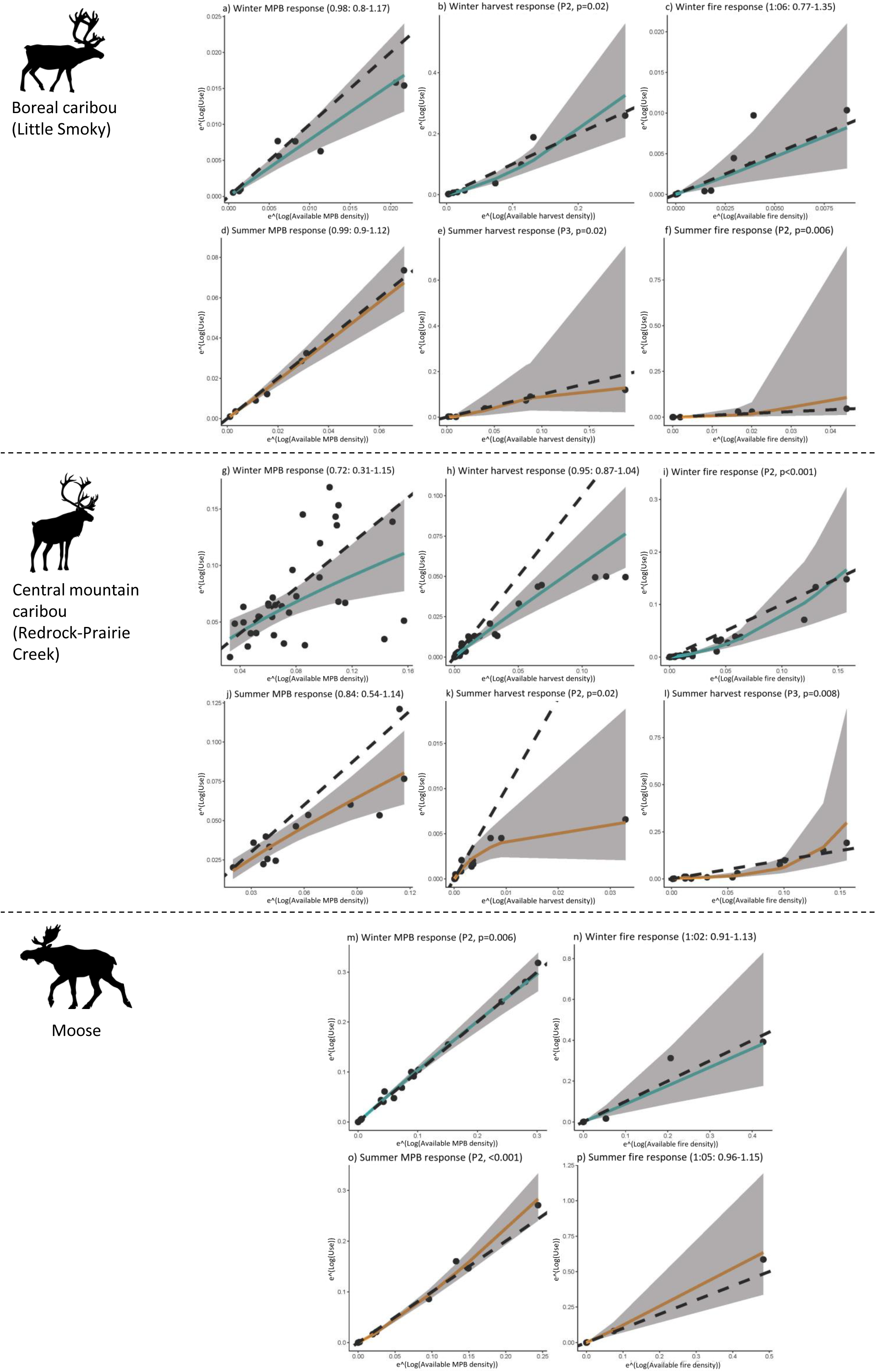
Functional responses in habitat use and selection by boreal caribou, central mountain caribou, and moose in west-central Alberta, Canada, during 2008-2010. Models included MPB, harvest, and fire density across both winter (blue) and summer (orange). Dashed lines indicate proportional habitat use and black dots indicate individuals. Shaded areas are 95% confidence intervals.

### Redrock-Prairie Creek (Central mountain caribou)

Central mountain caribou from the Redrock-Prairie Creek population avoided areas with higher densities of MPB during winter and selected areas with MPB during summer (Table 2, Fig. 3g&3j). These caribou avoided areas with higher densities of harvest and fire during winter and summer (Fig. 3h,i,k,l). They also selected open habitats, high elevations, and flatter slopes across winter and summer (Table 2). Both models had fair fit (winter=40.96%, summer=41.82%) and good to high predictive performance (mean winter r_s_=1, summer r_s_=0.75, S8-S9) (Mukaka, 2012).

**Table 2.** Estimated β coefficients, standard errors (SE), lower and upper 95% confidence intervals, Z-values, and p-values of central mountain caribou habitat selection model during winter and summer in west-central Alberta, Canada, during 2008-2010. Reference category for habitat was ‘forested’.

| Fixed effects | Estimate | SE | Lower 95% CI | Upper 95% CI | Z-value | p-value |
| --- | --- | --- | --- | --- | --- | --- |
| **Central mountain caribou: Winter** |  |  |  |  |  |  |
| Intercept | -2.870 | 0.034 | -2.936 | -2.804 | -85.079 | <0.001 |
| Habitat [Open] | 0.739 | 0.015 | 0.711 | 0.768 | 51.058 | <0.001 |
| Elevation (m) | -5.932 | 0.374 | -6.665 | -5.199 | -15.861 | <0.001 |
| Elevation<sup>2</sup> (m) | 6.434 | 0.374 | 5.700 | 7.167 | 17.187 | <0.001 |
| Slope | 0.691 | 0.026 | 0.641 | 0.742 | 26.850 | <0.001 |
| Slope<sup>2</sup> | -1.275 | 0.027 | -1.328 | -1.221 | -46.757 | <0.001 |
| Harvest | 1.146 | 0.028 | 1.090 | 1.201 | 40.452 | <0.001 |
| Harvest<sup>2</sup> | -2.295 | 0.049 | -2.390 | -2.199 | -47.062 | <0.001 |
| Fire | -0.120 | 0.018 | -0.155 | -0.084 | -6.663 | <0.001 |
| Fire<sup>2</sup> | -0.022 | 0.018 | -0.058 | 0.014 | -1.209 | 0.226 |
| MPB | 0.533 | 0.016 | 0.503 | 0.563 | 34.460 | <0.001 |
| MPB<sup>2</sup> | -0.458 | 0.016 | -0.489 | -0.426 | -28.408 | <0.001 |
| **Central mountain caribou: Summer** |  |  |  |  |  |  |
| Intercept | -3.028 | 0.103 | -3.229 | -2.827 | -29.492 | <0.001 |
| Habitat [Open] | 1.068 | 0.026 | 1.017 | 1.118 | 41.709 | <0.001 |
| Elevation (m) | 14.090 | 1.006 | 12.117 | 16.062 | 14.000 | <0.001 |
| Elevation<sup>2</sup> (m) | -13.318 | 0.996 | -15.270 | -11.366 | -13.375 | <0.001 |
| Slope | 0.943 | 0.052 | 0.841 | 1.046 | 18.044 | <0.001 |
| Slope<sup>2</sup> | -1.319 | 0.052 | -1.319 | -1.217 | -25.341 | <0.001 |
| Harvest | 1.057 | 0.060 | 0.939 | 1.175 | 17.594 | <0.001 |
| Harvest<sup>2</sup> | -1.832 | 0.139 | -2.104 | -1.560 | -13.196 | <0.001 |
| Fire | 0.298 | 0.031 | 0.238 | 0.358 | 9.750 | **<0.001** |
| Fire<sup>2</sup> | -0.285 | 0.032 | -0.347 | -0.223 | -8.995 | **<0.001** |
| MPB | -0.198 | 0.026 | -0.247 | -0.149 | -7.893 | **<0.001** |
| MPB<sup>2</sup> | 0.186 | 0.028 | 0.132 | 0.240 | 6.766 | **<0.001** |

FR models indicated that mountain caribou did not exhibit a functional response to MPB during winter, though there was a nonsignificant trend indicating avoidance in home ranges with higher MPB densities (Fig. 4g). During summer, they displayed a functional response, with avoidance increasing as MPB densities increased across home ranges (Fig. 4j). These caribou also displayed functional responses to harvest in both winter and summer, with greater avoidance as harvest densities increased across home ranges (Fig. 4h&4k). Finally, these mountain caribou displayed a functional response to fire in both winter and summer, avoiding fire at lower densities but with use increasing as fire densities increased across home ranges (Fig. 4i&4l).

### Moose

Moose generally selected areas with higher densities of MPB and fire during summer and winter (Table 3, Fig. 3m-p). Moose selected open habitat during winter, forested habitat during summer, lower elevations and flatter slopes during winter, and moderately higher elevations and flatter slopes during summer, though they rarely used highest elevations within their home ranges (Table 3). The winter and summer models had weak fit (winter=29.63%, summer=6.42%, S8).

**Table 3.** Estimated β coefficients, standard errors (SE), lower and upper 95% confidence intervals, Z-values, and p-values of moose habitat selection model during winter and summer in west-central Alberta, Canada, during 2008-2010. Reference category for habitat was ‘forested’.

| Fixed effects | Estimate | SE | Lower 95% CI | Upper 95% CI | Z-value | p-value |
| --- | --- | --- | --- | --- | --- | --- |
| **Moose: Winter** |  |  |  |  |  |  |
| Intercept | -2.150 | 0.234 | -2.608 | -1.691 | -9.189 | **<0.001** |
| Habitat [Open] | 0.192 | 0.024 | 0.146 | 0.239 | 8.073 | **<0.001** |
| Sex [Male] | -0.674 | 0.312 | -1.285 | -0.062 | -2.160 | **0.031** |
| Elevation (m) | 9.567 | 0.723 | 8.150 | 10.984 | 13.235 | **<0.001** |
| Elevation<sup>2</sup> (m) | -10.341 | 0.726 | -11.764 | -8.918 | -14.247 | **<0.001** |
| Slope | -0.591 | 0.035 | -0.659 | -0.523 | -16.954 | **<0.001** |
| Slope<sup>2</sup> | 0.357 | 0.038 | 0.282 | 0.432 | 9.313 | **<0.001** |
| Fire | -1.299 | 0.058 | -1.412 | -1.185 | -22.459 | **<0.001** |
| Fire<sup>2</sup> | 1.196 | 0.048 | 1.103 | 1.290 | 25.094 | **<0.001** |
| MPB | 0.204 | 0.054 | 0.097 | 0.311 | 3.746 | **<0.001** |
| MPB<sup>2</sup> | -0.100 | 0.047 | -0.192 | -0.008 | -2.128 | **0.033** |
| **Moose: Summer** |  |  |  |  |  |  |
| Intercept | -2.451 | 0.119 | -2.683 | -2.218 | -20.681 | **<0.001** |
| Habitat [Open] | -0.239 | 0.031 | -0.299 | -0.180 | -7.854 | **<0.001** |
| Sex [Male] | 0.213 | 0.167 | -0.114 | 0.540 | 1.278 | 0.201 |
| Elevation (m) | 1.850 | 0.920 | 0.047 | 3.653 | 2.011 | **0.044** |
| Elevation<sup>2</sup> (m) | -1.887 | 0.919 | -3.687 | -0.087 | -2.054 | **0.040** |
| Slope | 0.884 | 0.058 | 0.771 | 0.997 | 15.311 | **<0.001** |
| Slope<sup>2</sup> | -1.020 | 0.060 | -1.137 | -0.904 | -17.143 | **<0.001** |
| Fire | -0.111 | 0.077 | -0.262 | 0.040 | -1.442 | 0.149 |
| Fire<sup>2</sup> | -0.023 | 0.018 | 0.142 | 0.403 | -1.209 | **<0.001** |
| MPB | -0.890 | 0.072 | -1.031 | -0.748 | -12.350 | **<0.001** |
| MPB<sup>2</sup> | 0.787 | 0.056 | 0.676 | 0.897 | 13.962 | **<0.001** |

K-fold cross-validation demonstrated that both models had good predictive performance (mean winter r_s_=0.87, summer r_s_=0.82, S8-S9) (Mukaka, 2012). While non-linearity was detected in MPB FR models, in general moose use of MPB and fire was largely proportional to availability across home ranges (Fig. 4m-4p).

## Discussion

### Responses to MPB

As predicted, caribou avoided MPB during winter. Trends in FR models indicated that avoidance of MPB may increase with increasing densities of MPB across home ranges, but this was not statistically significant. In contrast with our predictions, trends were reversed during summer with caribou selecting areas with higher densities of MPB. However, our functional response models did indicate that, for mountain caribou, use was lower than expected given availability across home ranges. As predicted, we also found that moose selected areas with higher densities of MPB during both seasons.

Caribou display seasonal shifts in diet (i.e., lichen during winter, vascular plants during summer) which may be impacted by MPB-associated changes in understory vegetation (Webber *et al*., 2022). The avoidance of MPB we observed during winter may be due to loss of lichen cover soon after infestation (3-5 years; Cichowski *et al*., 2022). However, studies from the same area suggest lichen cover may not be impacted by MPB even 10-15 years post-infestation (Nobert et al., 2020). It is possible that the avoidance of MPB we observed is, therefore, driven by other environmental factors that have been impacted by MPB. For example, increased snow depth under MPB-thinned canopies may negatively impact caribou access and movement (Cichowski, 2010). Conversely, we found that moose used areas with MPB during winter, likely due to the associated increase in availability of vascular plants (Cichowski *et al*., 2022). Moose are a primary prey of wolves (James *et al*., 2004), so it is possible that moose use of MPB areas is driving increased wolf presence, which in turn is driving caribou avoidance of MPB (i.e., disturbance-mediated apparent competition, DMAC; Neufeld *et al*., 2021). In our study area, caribou are at increased mortality risk where they co-occur with moose (Peters, 2010).

The switch from avoidance to use of MPB-impacted areas by caribou during summer may be due to seasonal consumption of vascular plants in MPB-impacted areas (Webber *et al*., 2022). It is also possible that whatever factor is driving avoidance in winter is no longer limiting during summer. For example, snowmelt may make these areas accessible, and wolves preferentially prey on large ungulates during winter (Stahler, Smith and Guernsey, 2006). Summer use of MPB-impacted areas by caribou suggests it is not inaccessibility due to treefall driving avoidance during winter, as this would continue into summer (Lewis and Hartley, 2006a).

FR models indicated that MPB use was lower than expected given availability of MPB across mountain caribou summer home ranges. RSF models also showed that these caribou used open areas and higher elevations, in accordance with observed range contractions and movement to higher elevations year-round (MacNearney *et al*., 2016; Williams *et al*., 2021). The functional response we observed during summer may reflect the lower densities of MPB in alpine and subalpine areas (Sambaraju and Goodsman, 2021). However, with temperatures rising due to climate change, the presence of MPB-damaged trees is increasing at higher elevations (Gibson *et al*., 2008). As these mountain caribou avoided MPB during winter, increased MPB infestations at higher elevations may further reduce habitat for this population, making them more vulnerable to mortalities from predation or stochastic events (MacNearney *et al*., 2016; Williams *et al*., 2021)

### Responses to harvest and fire

We found that responses to our proxies for MPB management varied across species and populations. As predicted, mountain caribou in Redrock-Prairie Creek avoided areas with higher densities of harvest and fire, whereas moose used areas with higher densities of fire. This reflects the typical habitat specialisations and responses to disturbance displayed by these species (Finnegan *et al*., 2021; Johnson and Rea, 2023a; Seip and Jones, 2008; Wasser *et al*., 2011). It is likely that these mountain caribou avoided disturbance due to the risk of increased mortality and reduced lichen availability (Lochhead, Kleynhans and Muhly, 2022; Waterhouse, Armleder and Nemec, 2011). In contrast, moose eat a variety of plants that are often abundant after fire, likely promoting moose use of burned areas (Gasaway *et al*., 1989; Johnson and Rea, 2023b; Mumma *et al*., 2024). Moose use of burned areas has been documented in multiple studies and sites, however this relationship is complex, and moose use may decrease depending on burn severity, burn size, and plant regrowth rates (Brown *et al*., 2018; DeMars *et al*., 2019; Gasaway *et al*., 1989; Mumma *et al*., 2024). Harvest was dropped from moose analyses as it was strongly, negatively correlated with elevation. However, moose selected areas in low-mid elevation ranges, indicating that moose may have preferentially used areas with relatively higher harvest. This supports previous findings indicating that moose utilise harvest disturbance, albeit again depending on factors such as age, plant regrowth, and patch size (Johnson and Rea, 2023a; Thompson and Vukelich, 1981). However, it must be noted that, while disturbance may benefit moose in terms of plant availability for consumption, they are also reliant upon intact forest availability for thermoregulation, with access being limited in heavily disturbed environments (Laforge *et al*., 2016). Therefore, while moose may exhibit general preference for disturbance, as shown here, continued assessments of moose behaviour under different disturbance levels are required (Agnew *et al*., 2022).

Functional responses also varied between species, and the directionality of mountain caribou functional responses in Redrock-Prairie Creek differed among disturbances. Moose did not exhibit a functional response to fire, with degree of selection remaining proportional as availability changed across home ranges. However, these mountain caribou increasingly avoided harvest as it became more available across home ranges, as was anticipated based on the literature (Schaefer and Mahoney, 2007). This relationship was inverted for fire, with these caribou easing avoidance of burned areas as they became more available across home ranges. This may reflect an inability to avoid burned areas within their already restricted home ranges (density of fire was higher than density of harvest within home ranges for this population, S3, Fig. 2) (MacNearney *et al*., 2016).

Boreal caribou from the Little Smoky population displayed more complex and variable relationships with our proxies for MPB management than moose or our focal mountain caribou. These caribou did not respond to fire during winter, but selected it during summer. We note that this population (Little Smoky) has low densities of fire in its overall range (Fig. 2, S3, Russell, Pendlebury and Ronson, 2016). It is, therefore, possible that this selection may reflect a preference for forested habitats where fires also occur, rather than selection for fire itself.

Evaluating functional responses to fire relative to other characteristics of home ranges (e.g., % undisturbed, landcover type) might help to explain this result further, but was beyond the scope of this study. Contrasting with previous research and our focal mountain caribou population’s responses (Finnegan *et al*., 2021; Schaefer and Mahoney, 2007), we found that these boreal caribou used harvested areas during winter. However, our functional response models indicated that individuals with lower levels of harvest in their winter ranges avoided it. These boreal caribou also avoided harvest in summer. At the time of this study, the Little Smoky population had the highest level of anthropogenic disturbance of any boreal caribou population in Canada, with almost 90% of its range being disturbed (Semeniuk et al., 2012). Little Smoky caribou avoid harvest at the landscape scale (i.e., when selecting home ranges within their greater population range), but this effect is typically weakened at the home range scale, which we assessed (DeCesare et al., 2012). It is possible that seasonal dependency on old-growth pine forests for lichen during winter may be driving use of remaining mature forest stands within their home ranges, even if the surrounding 5km radius area is disturbed by harvest making it seem like they are selecting for the harvest itself. Our FR models support this explanation, as individuals with less harvest in their home ranges were able to avoid harvest. This result highlights the importance of applying functional response models when interpreting patterns of habitat selection relative to disturbance, particularly in highly disturbed landscapes such as the Little Smoky range. In comparison, seasonal shifts in diet, reproductive status, and use of different habitat types likely drove avoidance of harvested areas during summer. For example, caribou are not as reliant on lichen consumption in mature stands during summer months, and female caribou with young calves may reduce use of all heavily harvested areas to avoid predation pressure (Viejou *et al*., 2018).

It is possible that caribou and moose responses will vary with the intensity and severity of harvesting and fire. For example, we used harvest as a proxy for salvage logging and all fire disturbance, including wildfire, as a proxy for prescribed burning. The avoidance of harvest observed in mountain caribou may be an underestimate of the true impact of MPB salvage logging. This is because we included partial cuts in our measure of harvest whereas salvage logging often involves complete clearcutting, which could elicit more severe responses (Thorn *et al*., 2018). Response to fire was also negative for mountain caribou (Redrock-Prairie Creek), which were already in ranges with high fire disturbance, but not for boreal caribou (Little Smoky), which were in ranges with low fire disturbance. This indicates that small amounts of low intensity burns may not be overtly negative for caribou, but this may be dependent on the level of disturbance (both fire and otherwise) already present within individual ranges.

Prescribed burning may, therefore, not be advisable where it contributes to the compounding effects of disturbance, but useful in other circumstances, particularly where it is not as intensive in damage as wildfire, though further research into this is required.

### Limitations and further research

The limited GPS location data available during the initial stages of MPB infestation means that some of our estimated relationships in FR models were uncertain. More data collected across broader spatiotemporal scales might reveal more robust patterns, as would extending analyses to assess responses at the landscape scale; i.e., second order (Johnson, 1980). Additionally, as stated, we used simplified proxies for MPB salvage logging and prescribed burns. Larger GPS datasets may enable more complex and direct modelling of MPB management actions, for example by including the size and % canopy removal of disturbances or by directly comparing different harvesting and fire types. This would help to tease out the initial findings presented by this study further. The effects of harvest disturbance on moose also requires further study, including disentanglement from the effects of elevation. Additionally, we recommend that future studies assess the responses of other species, particularly other generalist ungulates and wolves, to further unravel the potential role of DMAC and impacts on the greater ecosystem (Curveira Santos *et al*., 2024). Finally, while we assessed two designatable units of caribou (i.e., boreal and central mountain) we only assessed a single population from each. We, therefore, recommend that future research also extend into other populations in order to determine the transferability of these results.

### Conclusions and management implications

MPB typically attacks mature forests, which caribou are ecologically dependent upon during winter. It is, therefore, concerning that our results show caribou avoiding these MPB-disturbed areas during this season. Conversely, moose selected MPB, suggesting infestations may increase habitat for moose. When considered together, it would appear that moose selection for MPB may have negative effects on caribou via DMAC, though wolf responses to MPB need to be explored to directly assess this. Whether DMAC is the driving force for caribou avoidance of MPB, or whether MPB is having a more direct effect on caribou, our results indicate that MPB is further limiting habitat availability for this species within already disturbed landscapes. Additionally, our result show that the two most prevalent forms of MPB management currently being applied, harvest and prescribed burns, may not be appropriate in caribou ranges and may even further contribute to the reduction of available habitat (McClelland *et al*., 2023; Steinke *et al*., 2020).

The link between disturbance and caribou population declines is undeniable (DeMars *et al*., 2023; Festa-Bianchet *et al*., 2011), and using harvest or large-scale, intensive burns within caribou ranges may have greater consequences for caribou than the MPB infestations themselves (Schmiegelow *et al*., 2000). Alternative actions must, therefore, be explored. For example, Little Smoky caribou showed positive responses to fire where fire occurrence was limited, indicating that investigation into the benefits or impacts of appropriate fire management activities such as small-scale prescribed burning and Indigenous fire stewardship (Christianson *et al*., 2022) should be explored. Notably, these activities may also increase forest heterogeneity, making areas less susceptible to future MPB infestations (Hoffman *et al*., 2022a; Hoffman *et al*., 2022b; Hood, Baker and Sala, 2016). Other fine-scale management, such as single-tree cut and burn, may also be an effective strategy that balances MPB management and the ecological needs of caribou and moose (Nobert *et al*., 2020), but requires further investigation and may not be feasible at larger scales.

It must be noted that these vulnerable ungulate populations responded to MPB very soon after initial infestations. Given the accelerating impacts of climate change and compounding pressures faced by these species in the Anthropocene, our study highlights the need for more effective detection, response, and management of MPB, with the aim of minimising impacts on species of conservations concern. Finally, selected management actions must not only consider ecological needs, but also local community needs and the economic impacts of MPB and management, with the ultimate goal of ensuring sustainable outcomes for both wildlife and humans (DeFries, Foley and Asner, 2004).

## Author contributions

LLG, LF & ACB conceived the idea and designed the methodology. JD and HL produced the spatial layers and generated maps. LLG led data analysis, supported by SC, VMP, LF & ACB. LLG led the writing of the manuscript, revised by LF & ACB. All authors contributed critically and gave final approval for publication.

## Acknowledgements

Funding for this research was provided by the Federal-Provincial Mountain Pine Beetle Partnership. ACB was supported by the Canada Research Chairs program. Caribou data used in this project were provided by the Government of Alberta, with a special thank you to Alberta Environment and Protected Areas’ Fish and Wildlife Stewardship Branch (caribou recovery program) for facilitating data sharing and providing comments on the manuscript. Thank you also to Mark Hebblewhite, for providing the moose data used in this project.

## Conflict of interest

The authors declare no conflict of interest.

## Data availability

Woodland caribou are listed as Threatened under federal legislation, are vulnerable to hunting and disturbance, and are the subject of multiple federal and provincial recovery programmes. As such, their GPS telemetry locations are kept confidential and any data sharing is made at the discretion of the Government of Alberta. Moose data used in this study are available on Movebank: https://www.movebank.org/cms/webapp?gwt_fragment=page=studies,path=study178994931.

## Ethics statement

All data used in this study had previously been collected for other projects, and appropriate permissions were obtained prior to use. Caribou collaring was performed under the Government of Alberta’s Animal Care Protocol No. 008, and further detail is available in Hervieux et al. 2013. Moose collaring was performed under the University of Montana Animal Care and Use Protocol 056-56MHECS-010207 and 059-09MHWB122109, Alberta Sustainable Resource Development licenses no. 21803, 27086, 27088, 27090 and Parks Canada permit JNP-2007-952, and further detail is available in Peters et al. 2013 and on Movebank (https://www.movebank.org/cms/webapp?gwt_fragment=page%3Dstudies%2Cpath%3Dstudy178994931).

## Notes

https://www.movebank.org/cms/webapp?gwt_fragment=page%3Dstudies%2Cpath%3Dstudy178994931

### Competing Interest Statement

The authors have declared no competing interest.

