## Supplementary Captions for "Vulnerable caribou and moose populations display contrasting responses to mountain pine beetle outbreaks and management"

S1: Additional details regarding data screening and cleaning for caribou and moose GPS collar data, collected in west-central Alberta (2008-2010), and the variables extracted for modelling.

S2: Information on GPS location datasets (subdivided by species/designatable unit and season) used to assess caribou and moose response to mountain pine beetle in west-central Alberta, Canada, between 2008 and 2010. Information includes the number of individuals (i.e., ID-season-year combination), total fixes within each dataset, and the lowest and highest number of fixes for individuals in each dataset (i.e., least and most observed individuals’ number of fixes). These locations were used to generate 95% kernel density estimates at the individual (third order) level.

S3: Variables used within RSF models to assess caribou and moose response to mountain pine beetle (MPB) and MPB management in west-central Alberta, Canada between 2008 and 2010. Variable type and range of values for each dataset and season (boreal caribou, central mountain caribou, moose) are shown.

S4: Pairwise Spearman’s correlation coefficients for the predictor variables considered for caribou and moose RSF models used to assess response to MPB and MPB management in west-central Alberta from 2008- 2010. Variable details are outlined in S5. Tests for winter and summer and 1km and 5km scales for disturbance densities are depicted. The variable for land cover type is denoted as “treed” and the sex of the animal is denoted as “anml_sx”.

S5: Model structure and AIC values for each RSF model used to assess caribou and moose response to MPB and MPB management in west-central Alberta, Canada, 2008-2010. The most parsimonous model is indicated in bold.

S6: Spatial variogram computed with the residuals from the final caribou and moose (both winter and summer) RSF models used to assess response to MPB and MPB management in west-central Alberta, 2008-2010. Note that there are no erroneous spikes or patterns, indicating no patterns of spatial autocorrelation for any RSFs.

S7: Functional response model structure, and associated p-values outputted by the likelihood ratio tests, used to assess caribou and moose responses to changing availabilities of disturbance in individual home ranges in west-central Alberta, Canada, 2008-2010. The final model selected for each season-species/designatable unit combination (i.e., dataset) is in bold based on p-value. All disturbance variables are at the 5km radius and were log_e_-transformed.

S8: Descriptions of model fit, with the percentage of variation explained, and results of K-Fold cross validations for boreal caribou, central mountain caribou, and moose RSF models (n=6), built using data collected in west-central Alberta, Canada, from 2008-2010. For each model, R(FE) describes the percentage variation explained by the fixed effects alone and R(FE+RE) describes the total variation explained when the random effect (individual variation) is included. The individual Spearman correlation values for each 5-fold cross validation is provided, along with the mean.

S9: Evaluation of our RSF models (divided by species/designatable unit and season, made to assess response to MPB and MPB management in west-central Alberta, Canada (2008-2010), using area-adjusted frequency of categories (bins) of scores from the RSF scores. The correlation is calculated between scores and area-adjusted frequencies for 80% of the data in a 5-fold cross-validation scheme (individual and mean Spearman correlation scores are available in Table S8).
