## Supplementary material for "Vulnerable caribou and moose populations display contrasting responses to mountain pine beetle outbreaks and management": S1

S1: Additional details regarding data screening and cleaning for caribou and moose GPS collar data, collected in west-central Alberta (2008-2010), and the variables extracted for modelling.

1. *Cleaning caribou and moose GPS data*

We filtered caribou location data to only include points collected from 2008-2010, the period 3-5 years after the entry of MPB in the area. Moose location data were collected between 2008 and 2010. Caribou location data underwent initial screening and cleaning by the Government of Alberta, including the removal of locations with DOP (dilution of precision) >12 (note: further information on DOP and number of satellites was not provided with the dataset, thus could not be used in any further cleaning). Any locations with missing latitudes or longitudes or obviously erroneous locations (i.e., large movements in a short period of time or locations occurring after animal death/collar drop-off) were also removed prior to data sharing. Setting 4 hours as the minimum fix-rate, we removed individuals with a fix-rate of <90% (Frair *et al.*, 2010; Hebblewhite, Percy and Merrill, 2007). We set 2 hours as our maximum fix-rate, which was the maximum available for moose data. As several individuals in the caribou datasets had fixes every 1.5 hours, we down-sampled these individuals so that they fell within our maximum and minimum thresholds (i.e., we only retained locations with intervals between 2-4 hours). Next, we removed remaining inaccurate locations by filtering out GPS points with an incoming and outgoing speed that fell above the 99^th^ percentile (i.e., erroneous “spikes”) (Gupte *et al.*, 2022; Bjørneraas *et al.*, 2010). Finally, we checked the data for abnormal turning angles, which did not indicate the need for any further filtering (Gupte *et al.*, 2022).

For moose data, we removed individuals with a fix success rate <90%, with 4-hour intervals set as the minimum fix-rate (Frair *et al.*, 2010; Hebblewhite, Percy and Merrill, 2007), and points with speeds above the 99% percentile (Gupte *et al.*, 2022; Bjørneraas *et al.*, 2010), as described above. Again, we assessed the data for abnormalities in turning angle and determined that moose data did not require further filtering based on movement. We also removed 2D points (i.e., < 4 satellites) with a hDOP > 5 (Lewis *et al.*, 2007).

2. *Generating MPB rasters*

We used annual MPB survey data available from the Government of Alberta for the years of our study (2008-2010) and all previous years of MPB data (put in the year) (Government of Alberta, 2023). Datasets were derived from annual aerial surveys performed as part of ongoing forest health assessments across the province. Surveys were typically performed between late June and early August and followed strict collection protocols (Government of Alberta and ESRD, 2012; Government of Alberta, 2020). Surveyors noted the number and locations of new red stage (needle death) trees in each assigned area. These were saved as point locations unless they covered an area greater than 10ha, with these larger areas being saved as polygons (Government of Alberta and ESRD, 2012). Grey stage trees and red trees that had previously been documented were not captured. Red tree patches smaller than approximately three trees were also not captured. Notably, the survey areas selected each year were usually different, with the goal of covering a larger overall area over time (Government of Alberta, 2023). Therefore, many areas may only be surveyed every couple of years, with MPB often continuing to spread to neighbouring trees in the interim period. This, coupled with the fact that only new red stage trees are documented, means that the exact number and locations of trees impacted by MPB are likely underestimated in these datasets.

To provide a more accurate depiction of the true extent of MPB infestations, we combined point and polygon data on MPB occurrence for each year (i.e., each year of study with all previous years) and converted these point data into rasters (30X30m cells). If fire and harvest had disturbed >50% of a cell that year, or in any previous years (sources: Government of Alberta, 2022; ABMI, 2021), then we considered MPB at that cell to have been removed by these disturbances. Remaining cells with MPB occurrence were then converted back to points (Fig. 2a in the main text). As survey guidelines outlined that any counts of trees impacted by MPB in that area were likely to be an underestimate, each point represented a guaranteed occurrence of MPB in general at that 30x30m site, while disregarding the number of trees documented. We then used focal statistics to extract the density of MPB at 1km and 5km radii for each of the caribou and moose used and available GPS locations. These two scales are commonly used for these large ungulates (McKay and Finnegan, 2023; Leblond, Dussault and Ouellet, 2013) and are based on estimates of average daily movement (Bergman, Schaefer and Luttich, 2000; Danks and Porter, 2010).

3. *Additional Variables*

The National Terrestrial Ecosystem Monitoring System (NTEMS) is a satellite-data-driven series of information products created to aid in monitoring Canada’s forested ecosystems (White *et al.*, 2014). Using Landsat data, supplemented with LiDAR data and ground plots (Mulverhill *et al.*, 2022; Wulder *et al.*, 2024), NTEMS consists of a series of forest information products, including year and type of forest disturbance (Hermosilla *et al.*, 2018; Hermosilla *et al.*, 2016). We used these high-resolution (30x30) raster layers for harvest and fire variables. We were primarily interested in wildlife responses to recent forest disturbances, so we used harvest and fires that occurred in the 20 years prior to each GPS location date (Lacerte, Leblond and St‐Laurent, 2021; Vors *et al.*, 2007; Finnegan *et al.*, 2021; Silva, 2020). Salvage harvesting in response to MPB infestations typically involves clearcutting impacted areas, but greater percentage tree retention has also been recommended and applied (Peter and Bogdanski, 2010). The intensity of the harvesting applied is often dependent upon the scale and intensity of the attack, as well as federal and provincial government policies (Peter and Bogdanski, 2010; ASRD, 2007). Therefore, we retained all harvest as a proxy for general MPB-associated harvest, whether it was used explicitly for this purpose or not (Fig. 2b in the main text). We used this information to produce harvest density variables at a 1km and 5km radius around each point (as outlined above). We also generated fire (burn) density variables at a 1km and 5km radius around each GPS location. Fire data included information predominantly on wildfire occurrences, although any fires are captured including prescribed burns. We considered this dataset as a proxy for fire disturbance from prescribed burning for our analysis (Hermosilla *et al.*, 2016) (Fig. 2c in the main text).

Additional variables extracted for modelling included landcover type (forested or open), elevation, and slope. These variables were selected as they have previously been shown to influence species’ habitat selection (Hebblewhite *et al.*, 2010; DeCesare *et al.*, 2012; Peters, 2010). For land cover type, we used NTEMS land cover layers for the same years as our GPS data (2008-2010) (Hermosilla *et al.*, 2016). Elevation and slope were extracted using a 30x30m digital elevation model (Natural Resources Canada, 2011). Further details on variables are available in Table S3.
