## Supplementary material for "Vulnerable caribou and moose populations display contrasting responses to mountain pine beetle outbreaks and management": S2

S2. Information on GPS location datasets (subdivided by species/designatable unit and season) used to assess caribou and moose response to mountain pine beetle in west-central Alberta, Canada, between 2008 and 2010. Information includes the number of individuals (i.e., ID-season-year combination), total fixes within each dataset, and the lowest and highest number of fixes for individuals in each dataset (i.e., least and most observed individuals’ number of fixes). These locations were used to generate 95% kernel density estimates at the individual (third order) level.

| Dataset | #Individuals  (ID-Season-Year) | Total used locations | Min. used locations/individual | Max. used locations/individual |
| --- | --- | --- | --- | --- |
| Boreal caribou (Winter) | 9 | 11,487 | 241 | 2,447 |
| Boreal caribou (Summer) | 7 | 6,929 | 479 | 1,645 |
| Central mountain caribou (Winter) | 32 | 46,701 | 354 | 2,474 |
| Central mountain caribou (Summer) | 13 | 13,520 | 244 | 1,557 |
| Moose (Winter) | 16 | 14,826 | 289 | 1,739 |
| Moose (Summer) | 8 | 10,266 | 865 | 1,760 |
