## Supplementary material for "Vulnerable caribou and moose populations display contrasting responses to mountain pine beetle outbreaks and management": S3

S3: Variables used within RSF models to assess caribou and moose response to mountain pine beetle (MPB) and MPB management in west-central Alberta, Canada between 2008 and 2010. Variable type and range of values for each dataset and season (boreal caribou, central mountain caribou, moose) are shown.

|  |  | Boreal caribou | | Central mountain caribou | | Moose | |
| --- | --- | --- | --- | --- | --- | --- | --- |
| Variable | Description | Winter | Summer | Winter | Summer | Winter | Summer |
| Land cover type | forested^1^ or  open^2^; categorical. | NA | NA | NA | NA | NA | NA |
| Elevation | Metres above mean sea level. | 993-1734 | 976-1377 | 886-2628 | 1040-2544 | 733-2438 | 730-2435 |
| Slope | Slope in degrees. | 0-26.8 | 0-26.6 | 0-72.8 | 0-70.6 | 0-68.8 | 0-60.83 |
| MPB (1km) | Density estimate of MPB disturbance within 1km. | 0-0.36 | 0-0.24 | 0-0.74 | 0-0.74 | 0-0.70 | 0-0.72 |
| MPB (5km) | Density estimate of MPB disturbance within 5km. | 0-0.13 | 0-0.1 | 0-0.44 | 0-0.44 | 0-0.44 | 0-0.44 |
| Harvest (1km) | Density estimate of harvest disturbance within 1km. | 0-0.82 | 0-0.76 | 0-0.91 | 0-0.83 | NA | NA |
| Harvest (5km) | Density estimate of harvest disturbance within 5km. | 0-0.42 | 0-0.44 | 0-0.45 | 0-0.26 | NA | NA |
| Fire (1km) | Density estimate of fire disturbance within 1km. | 0-0.71 | 0-0.72 | 0-0.99 | 0-0.99 | 0-0.99 | 0-0.99 |
| Fire (5km) | Density estimate of fire disturbance within 5km. | 0-0.05 | 0-0.05 | 0-0.77 | 0-0.78 | 0-0.78 | 0-0.77 |
| Sex | Male or female. | - | - | - | - | M or F | M or F |
