## Supplementary figures and images for "Vulnerable caribou and moose populations display contrasting responses to mountain pine beetle outbreaks and management"

### S4

#### Slide 1
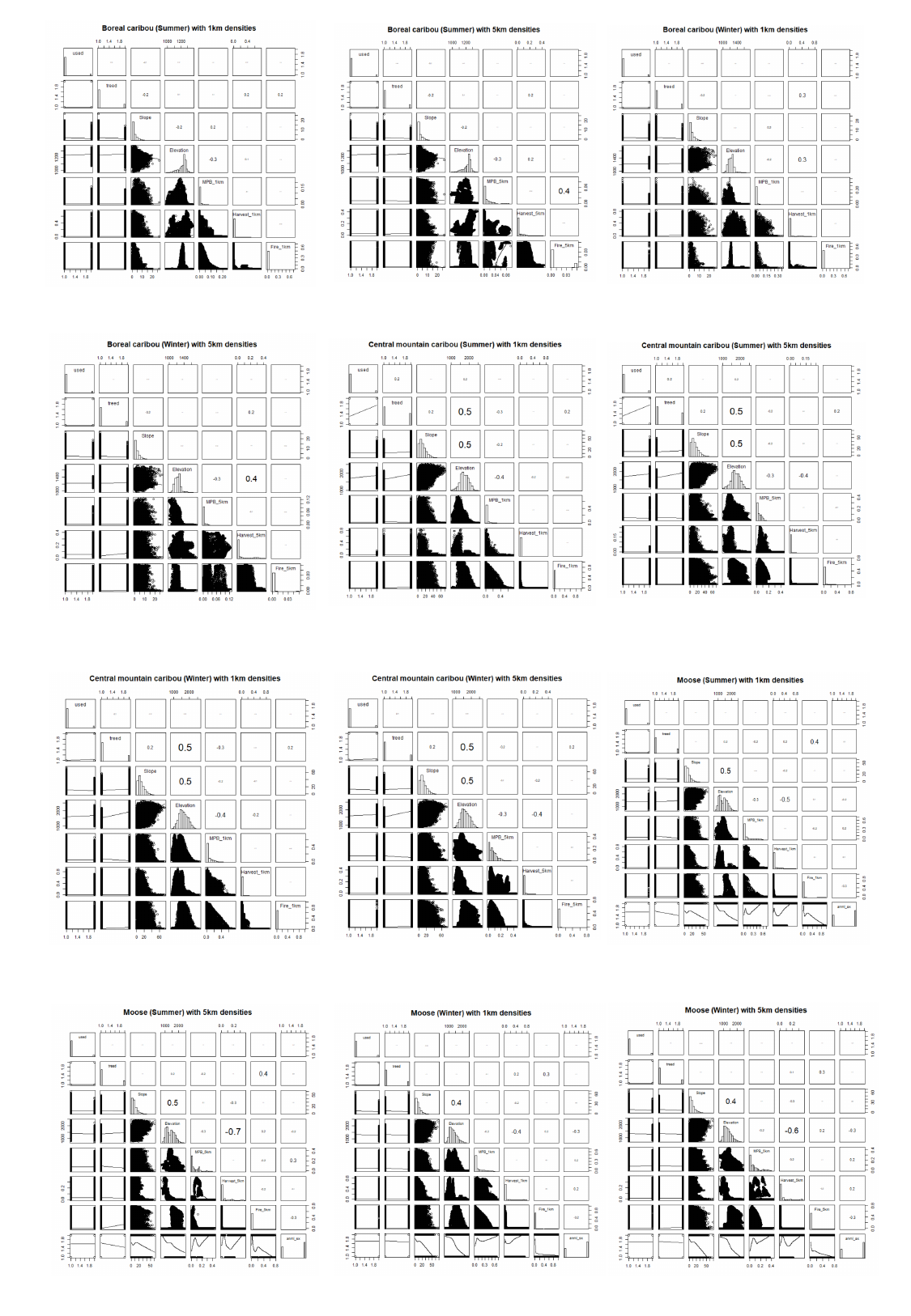

### S6

#### Slide 1
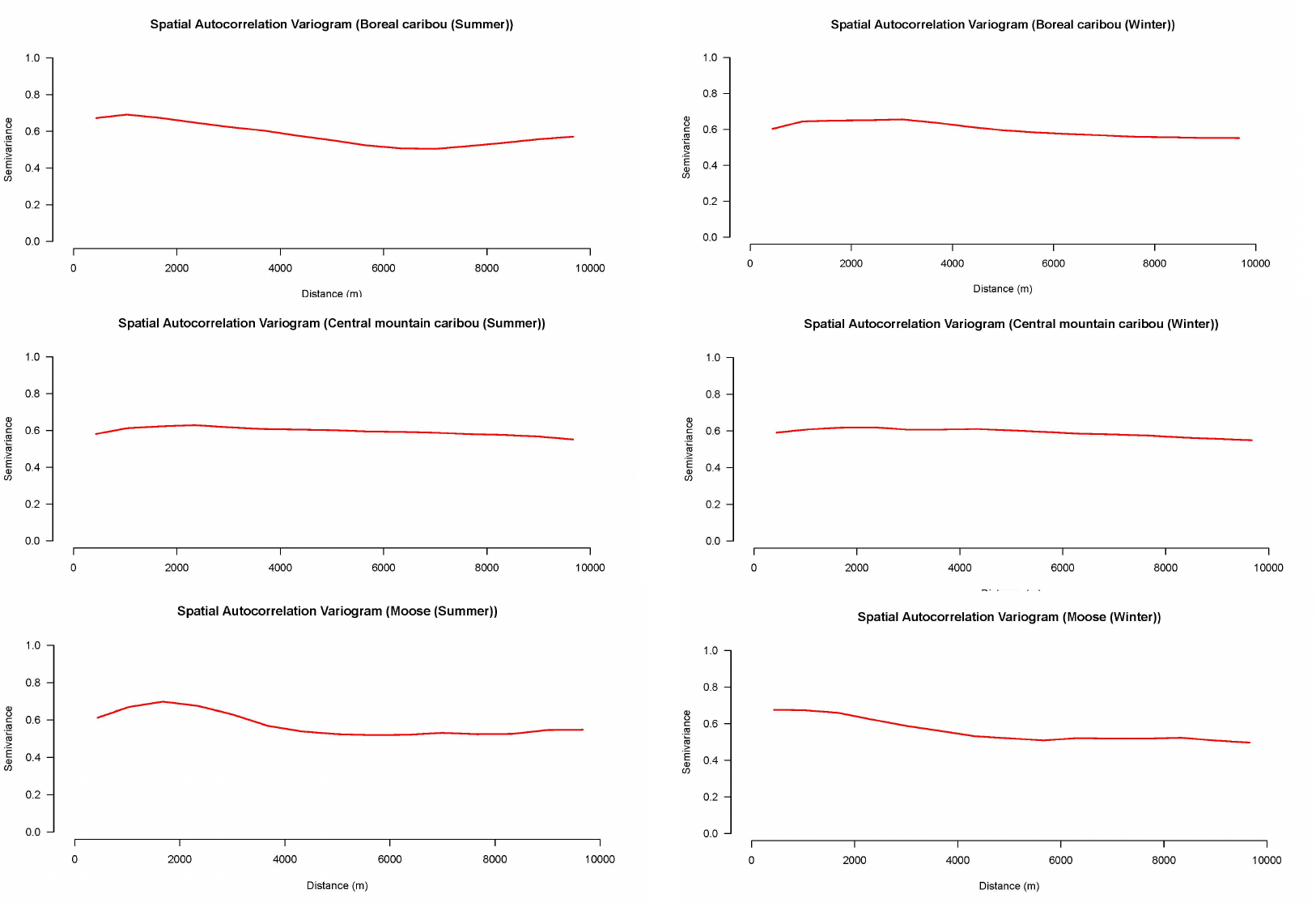

### S9

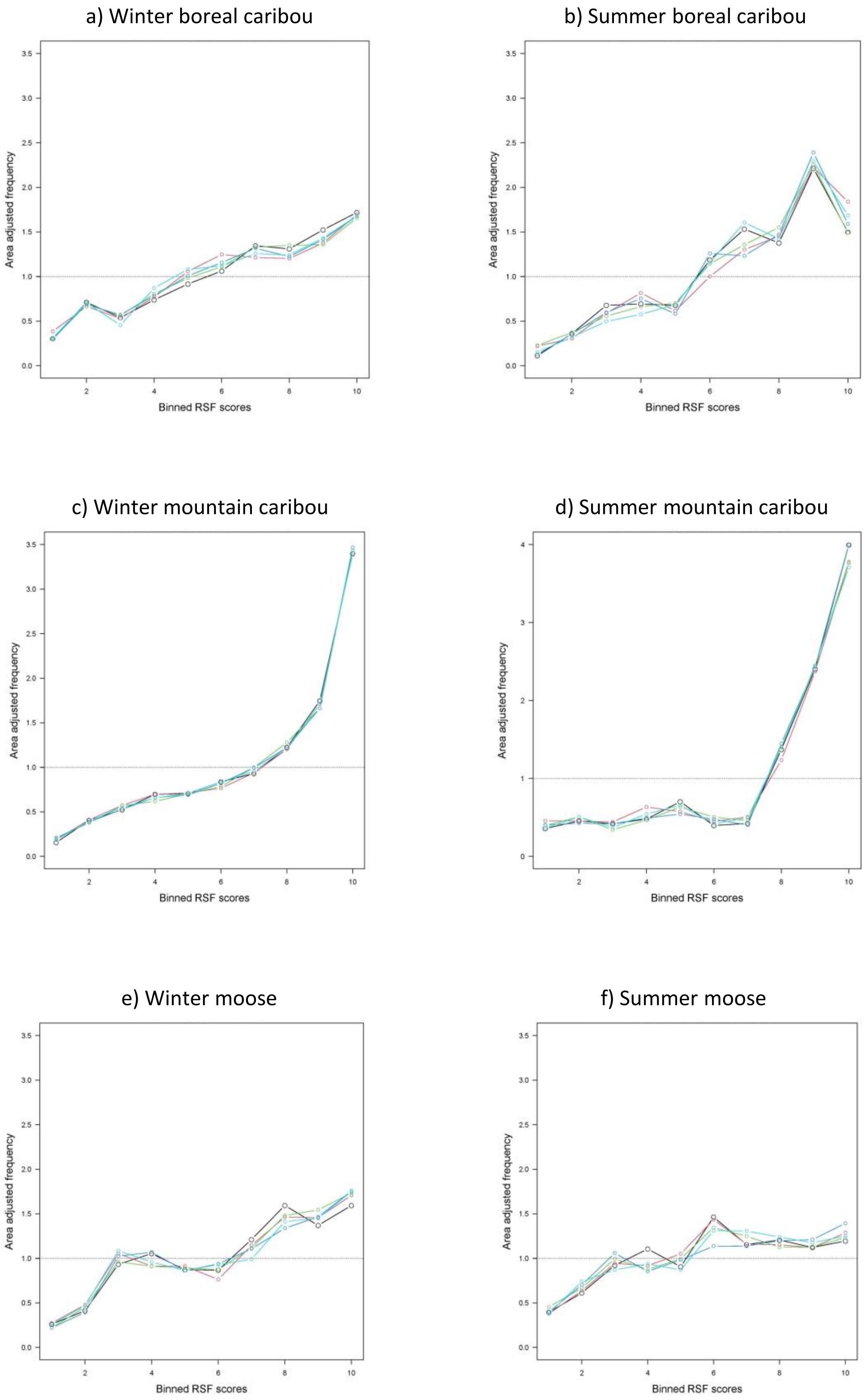
