## Supplementary material for "Vulnerable caribou and moose populations display contrasting responses to mountain pine beetle outbreaks and management": S5

S5: Model structure and AIC values for each RSF model used to assess caribou and moose response to MPB and MPB management in west-central Alberta, Canada, 2008-2010. The most parsimonous model is indicated in bold.

| Dataset | Model Structure | AIC |
| --- | --- | --- |
| Boreal caribou (winter; 1km) | Used(1/0) ~ land cover type + elevation + elevation^2^ + slope + slope^2^ + Harvest(1km) + Harvest(1km)^2^ + Fire(1km) + Fire(1km)^2^ + MPB(1km) + MPB(1km)^2^ + (1\|ID) | 74462.35 |
| Boreal caribou (winter; 5km) | Used(1/0) ~ land cover type + elevation + elevation^2^ + slope + slope^2^ + Harvest(5km) + Harvest(5km)^2^ + Fire(5km) + Fire(5km)^2^ + MPB(5km) + MPB(5km)^2^ + (1\|ID) | **74239.79** |
| Boreal caribou (summer; 1km) | Used(1/0) ~ land cover type + elevation + elevation^2^ + slope + slope^2^ + Harvest(1km) + Harvest(1km)^2^ + Fire(1km) + Fire(1km)^2^ + MPB(1km) + MPB(1km)^2^ + (1\|ID) | 42765.66 |
| Boreal caribou (summer, 5km) | Used(1/0) ~ land cover type + elevation + elevation^2^ + slope + slope^2^ + Harvest(5km) + Harvest(5km)^2^ + Fire(5km) + Fire(5km)^2^ + MPB(5km) + MPB(5km)^2^ + (1\|ID) | **42179.85** |
| Central mountain caribou (winter; 1km) | Used(1/0) ~ land cover type + elevation + elevation^2^ + slope + slope^2^ + Harvest(1km) + Harvest(1km)^2^ + Fire(1km) + Fire(1km)^2^ + MPB(1km) + MPB(1km)^2^ + (1\|ID) | 288281.50 |
| Central mountain caribou (winter; 5km) | Used(1/0) ~ land cover type + elevation + elevation^2^ + slope + slope^2^ + Harvest(5km) + Harvest(5km)^2^ + Fire(5km) + Fire(5km)^2^ + MPB(5km) + MPB(5km)^2^ + (1\|ID) | **287715.10** |
| Central mountain caribou (summer; 1km) | Used(1/0) ~ land cover type + elevation + elevation^2^ + slope + slope^2^ + Harvest(1km) + Harvest(1km)^2^ + Fire(1km) + Fire(1km)^2^ + MPB(1km) + MPB(1km)^2^ + (1\|ID) | **81009.83** |
| Central mountain caribou (summer; 5km) | Used(1/0) ~ land cover type + elevation + elevation^2^ + slope + slope^2^ + Harvest(5km) + Harvest(5km)^2^ + Fire(5km) + Fire(5km)^2^ + MPB(5km) + MPB(5km)^2^ + (1\|ID) | 81617.37 |
| Moose (winter; 1km) | Used(1/0) ~ land cover type + sex + elevation + elevation^2^ + slope + slope^2^ + Fire(1km) + Fire(1km)^2^ + MPB(1km) + MPB(1km)^2^ + (1\|ID) | 94977.65 |
| Moose (winter; 5km) | Used(1/0) ~ land cover type + sex + elevation + elevation^2^ + slope + slope^2^ + Fire(5km) + Fire(5km)^2^ + MPB(5km) + MPB(5km)^2^ + (1\|ID) | **94225.28** |
| Moose (summer; 1km) | Used(1/0) ~ land cover type + sex + elevation + elevation^2^ + slope + slope^2^ + Fire(1km) + Fire(1km)^2^ + MPB(1km) + MPB(1km)^2^ + (1\|ID) | 67982.94 |
| Moose (summer; 5km) | Used(1/0) ~ land cover type + sex + elevation + elevation^2^ + slope + slope^2^ + Fire(5km) + Fire(5km)^2^ + MPB(5km) + MPB(5km)^2^ + (1\|ID) | **67939.33** |
