## Supplementary material for "Vulnerable caribou and moose populations display contrasting responses to mountain pine beetle outbreaks and management": S7

S7: Functional response model structure, and associated p-values outputted by the likelihood ratio tests, used to assess caribou and moose responses to changing availabilities of disturbance in individual home ranges in west-central Alberta, Canada, 2008-2010. The final model selected for each season-species/designatable unit combination (i.e., dataset) is in bold based on p-value. All disturbance variables are at the 5km radius and were log_e_-transformed.

| Dataset | Season | Model Structure |  | Pr(>Chisq) |  |
| --- | --- | --- | --- | --- | --- |
|  |  |  | MPB | Harvest | Fire |
| Boreal caribou | Winter | 1) Mean_used ~ Mean_available 2) Mean_used ~ Mean_available + (Mean_available)^2^ 3) Mean_used ~ Mean_available + (Mean_available)^2^ + (Mean_available)^3^ | **[ref]**  0.736  0.077 | [ref]  **0.023**  0.632 | **[ref]**  0.173  0.452 |
| Boreal caribou | Summer | 1) Mean_used ~ Mean_available 2) Mean_used ~ Mean_available + (Mean_available)^2^ 3) Mean_used ~ Mean_available + (Mean_available)^2^ + (Mean_available)^3^ | **[ref]**  0.227  0.155 | [ref]  0.139  **0.021** | [ref]  **0.006**  0.837 |
| Mountain caribou | Winter | 1) Mean_used ~ Mean_available 2) Mean_used ~ Mean_available + (Mean_available)^2^ 3) Mean_used ~ Mean_available + (Mean_available)^2^ + (Mean_available)^3^ | **[ref]**  0.096  0.155 | **[ref]**  0.243  0.341 | [ref]  **<0.001**  0.699 |
| Mountain caribou | Summer | 1) Mean_used ~ Mean_available 2) Mean_used ~ Mean_available + (Mean_available)^2^ 3) Mean_used ~ Mean_available + (Mean_available)^2^ + (Mean_available)^3^ | **[ref]**  0.341  0.825 | [ref]  **0.021**  0.554 | [ref]  0.344  **0.008** |
| Moose | Winter | 1) Mean_used ~ Mean_available 2) Mean_used ~ Mean_available + (Mean_available)^2^ 3) Mean_used ~ Mean_available + (Mean_available)^2^ + (Mean_available)^3^ | [ref]  **0.006**  0.525 | NA | **[ref]**  0.808  0.171 |
| Moose | Summer | 1) Mean_used ~ Mean_available 2) Mean_used ~ Mean_available + (Mean_available)^2^ 3) Mean_used ~ Mean_available + (Mean_available)^2^ + (Mean_available)^3^ | [ref]  **<0.001**  0.147 | NA | **[ref]**  0.101  0.133 |
