## Supplementary material for "Vulnerable caribou and moose populations display contrasting responses to mountain pine beetle outbreaks and management": S8

Table S8: Descriptions of model fit, with the percentage of variation explained, and results of K-Fold cross validations for boreal caribou, central mountain caribou, and moose RSF models (n=6), built using data collected in west-central Alberta, Canada, from 2008-2010. For each model, R(FE) describes the percentage variation explained by the fixed effects alone and R(FE+RE) describes the total variation explained when the random effect (individual variation) is included. The individual Spearman correlation values for each 5-fold cross validation is provided, along with the mean.

| Model | R(FE) | R(FE+RE) | Individual Spearman correlation (r, 5-fold) | Mean Spearman correlation (r) |
| --- | --- | --- | --- | --- |
| Boreal caribou: Winter | 20.12% | 23.53% | 0.98, 0.94, 0.99, 0.98, 0.98 | 0.97 |
| Boreal caribou: Summer | 30.78% | 51.81% | 0.92, 0.98, 0.96, 0.94, 0.98 | 0.96 |
| Central mountain caribou: Winter | 40.35% | 40.96% | 1, 1, 1, 1, 1 | 1 |
| Central mountain caribou: Summer | 39.48% | 41.82% | 0.75, 0.74, 0.75, 0.74, 0.78 | 0.75 |
| Moose: Winter | 21.33% | 29.63% | 0.87, 0.87, 0.88, 0.90, 0.84 | 0.87 |
| Moose: Summer | 4.50% | 6.42% | 0.78, 0.82, 0.77, 0.96, 0.77 | 0.82 |
